# Mesocorticolimbic reinforcement learning of reward representation and value provides an integrated mechanistic account for schizophrenia

**DOI:** 10.1101/2025.08.10.669522

**Authors:** Kenji Morita, Arvind Kumar

## Abstract

Mesocorticolimbic dopamine projections are crucial for value learning, motivational control, and cognitive functions, but their precise neurocomputational roles remain elusive. Based on recent experimental and theoretical findings, we constructed a neural circuit model where dopamine neuronal populations receive differential inputs from individual rewards and encode heterogeneous reward prediction errors, which train cortical and striatal neurons to learn reward-associated state representation and value. Learning is achieved via simultaneous ‘alignments’ of the cortical and striatal downstream connections to the mesocorticolimbic dopamine projections, and inhibition-dominance in the cortical recurrent network is a key for successful learning. Excessive excitation, whether pre-existing or induced by manipulations, leads to aberrant activity, which disrupts the alignments and even causes anti-alignment. This impairs both reward-specific motivational control and credit assignment, potentially explaining the negative and positive symptoms of schizophrenia, respectively. Our model thus provides a mechanistic account for schizophrenia, integrating the different causes and symptoms, with testable predictions.

## Introduction

Mesocorticolimbic dopamine (DA) has been suggested to be crucial for value learning, motivational control, and cognitive functions. Among them, the role of mesolimbic DA in value learning has largely been established under the reinforcement learning (RL) framework: DA encodes reward-prediction-error (RPE) and its modulation of cortico-striatal plasticity implements RPE-dependent update of value prediction ^1-3^ (although further consideration continues ^4-9^). Compared to this, DA’s roles in motivational and cognitive controls remain more elusive. A key feature of motivational control is its specificity to reward identity - seeking food when hungry and drink when thirsty, and reward identity-dependence of DA signals has been demonstrated ^10-12^. On the contrary, the conventional RL describes reward as one-dimensional scalar variable, ignoring reward identity/diversity. Regarding DA’s cognitive functions, early studies focused on how it modulates working memory ^13^, but DA also relates to wider prefrontal functions ^14, 15^, including the learning of sensorimotor associations ^16^ and the encoding of context representation ^17^. Whether these DA’s motivational and cognitive functions can also be understood in the RL framework remains elusive.

Psychiatric disorders such as schizophrenia provide a further window into the diverse computational roles of DA. DA receptor agonists and antagonists have psychomimetic and antipsychotic effects, respectively ^18, 19^. Psychosis, including delusion and hallucination, is a major aspect of schizophrenia, named the positive symptoms, and in the RL framework, can be seen as an issue in credit assignment (c.f., ^20^). Schizophrenia also has other major aspects, negative symptoms and cognitive impairments ^21, 22^, including motivational impairments such as avolition and anhedonia ^23^ and deficits in value representation ^24^. Previous studies have shown that certain aspects or symptoms of schizophrenia are associated with particular impairments in RL tasks ^25, 26^, including impaired goal-directed RL vs spared simple RL in stable environments ^24, 27^ and impairments coming from working memory deficits ^28-30^. However, RL-based integrated understanding of the diverse symptoms has yet to be obtained, possibly mirroring the lack of RL-based understanding of normal motivational and cognitive functions. Also, DA, glutamate, and sociodevelopmental effects were suggested as major factors of schizophrenia ^31, 32^, and they are proposed to converge onto changes in the cortical excitation/inhibition (E/I) balance ^32-35^, but its relation to RL remains unclear.

Recently, advanced RL models have been developed to account for either the motivational or cognitive function of DA, separately. Regarding the motivational function, the reward bases (RB) model ^36^ was developed in reference to reward identity-dependence of DA signals ^10-12^ and previous models ^37-40^. RB incorporates multi-dimensional RPE encoding by DA neuronal populations, which enables reward-specific motivational control. Regarding the cognitive function, the online value-Recurrent-Neural-Network (OVRNN) ^41^ was developed by elaborating the original value-RNN ^42^ to incorporate biologically plausible online local feedback. OVRNN enables learning of task-appropriate state representation and value by training of cortical recurrent-neural-network (RNN) and its readout (striatum) by RPE. RB does not deal with cortical DA’s functions, while OVRNN sticks to one-dimensional RPE, and so integration of these two models is desired. However, operation of RB is based on the ‘alignment’ of the striatum-DA connections to the backward mesolimbic connections, whereas OVRNN is based on the ‘alignment’ of the corticostriatal connections to the mesocortical connections, and whether both alignments can simultaneously occur is nontrivial. Moreover, even if they occur, it is not clear whether an integrated model can further explain the converged involvement of E/I imbalance in schizophrenia.

In the present study, we successfully addressed these issues. We developed a model of the mesocorticolimbic system that combined the RB and OVRNN models, and demonstrated that the two alignments can simultaneously occur. Crucially, we found that this occurrence of alignments depends on the E/I balance of the cortical recurrent RNN. In excitation-dominant regime, whether from the beginning or by a later balance shift or learning-rate bias, there appears aberrant persistent activity, which degrades or even reverses the alignments, resulting in impairments of both motivational control and credit assignment. Thus, we provide an instance of how the E/I-balance can affect computations besides controlling the stability of brain dynamics, and through that our model provides an integrated account of the diverse causes and symptoms of schizophrenia.

## Results

### Online-Value-RNN-Reward-Bases (OVRNN-RB) model

We constructed a model of the mesocorticolimbic system, by integrating two recent models: the online value-RNN (OVRNN) ^41^ and the reward bases (RB) ^36^. OVRNN, a biologically plausible version of the original value-RNN ^8, 42^, consists of an RNN (corresponding to cortex), a readout unit (striatum), and an error unit (DA neurons). The error unit calculates a scalar (i.e., one-dimensional) temporal-difference reward-prediction-error (TD-RPE), which is sent to the striatal readout and also to the cortical RNN via fixed random weights. In the RNN-readout (cortico-striatal) weights, state value is learned, as in the standard reinforcement learning (RL) model of basal ganglia ^43^. In the meantime, in the cortical RNN, state representation appropriate for the current task is learned, by virtue of an "alignment" (c.f., ^44-46^) of the RNN-readout (cortico-striatal) weights to the fixed random error-feedback (mesocortical) weights.

On the other hand, RB ^36^, developed in reference to previous studies ^37-40^, consists of cortical inputs, striatal value units, and DA error units. The DA units are heterogeneous as they receive differential inputs from multiple different rewards, encoding multi-dimensional TD-RPEs as a whole. The TD-RPEs are sent to the striatal units via fixed random weights, and used for the training of cortico-striatal weights, as well as the training of striatum-DA weights. Through learning, an "alignment" of the striatum-DA weights to the fixed random DA-striatum weights occurs: for instance, if a striatal unit receives a strong feedback from a DA unit that is strongly activated by a particular reward, the forward connection from this striatal unit to this DA unit becomes also strong. This alignment enables reward-specific value encoding and motivational control. Specifically, when that particular reward is highly desired (e.g., food when hungry), raised tonic DA (c.f., ^37, 39, 47^) from that DA unit can amplify, via physiological modulation, the input from the "corresponding" striatal unit, resulting in a specific amplification of the value of that reward ^36^.

As such, OVRNN deals with cortical RNN dynamics and its training by TD-RPE but not heterogeneity of rewards and TD-RPE signals, whereas RB deals with the latter but not the former. We constructed a model, OVRNN-RB (Fig. 1A,B), which combined OVRNN and RB, incorporating both heterogeneous TD-RPEs and training of cortical RNN by them. As mentioned above, learning of OVRNN is ensured by the alignment of the RNN-downstream weights to the DA-RNN (mesocortical) weights, while learning of RB is achieved through the alignment of the striatum-DA weights to the DA-striatum (mesolimbic) weights. When OVRNN and RB are combined, whether these two alignments can both occur is nontrivial, especially given that the RNN-downstream weights become more complex than those in OVRNN because there are now multiple striatal and DA units. As in the biologically most plausible version of OVRNN ("oVRNNrf-bio" in ^41^), we imposed biological constraints that the activity of RNN units and the weights of the RNN-striatum, striatum-DA, DA-RNN, and DA-striatum connections were non-negative and also the dependence of the update (plasticity) of the RNN weights was monotonic(+saturation). In addition, because it was shown that in OVRNN the mean of the weights onto the RNN units became negative (i.e., inhibition-dominance in the E/I balance) through learning ^41^, here we initialized the RNN weights to be negative on average.

**Figure 1.**
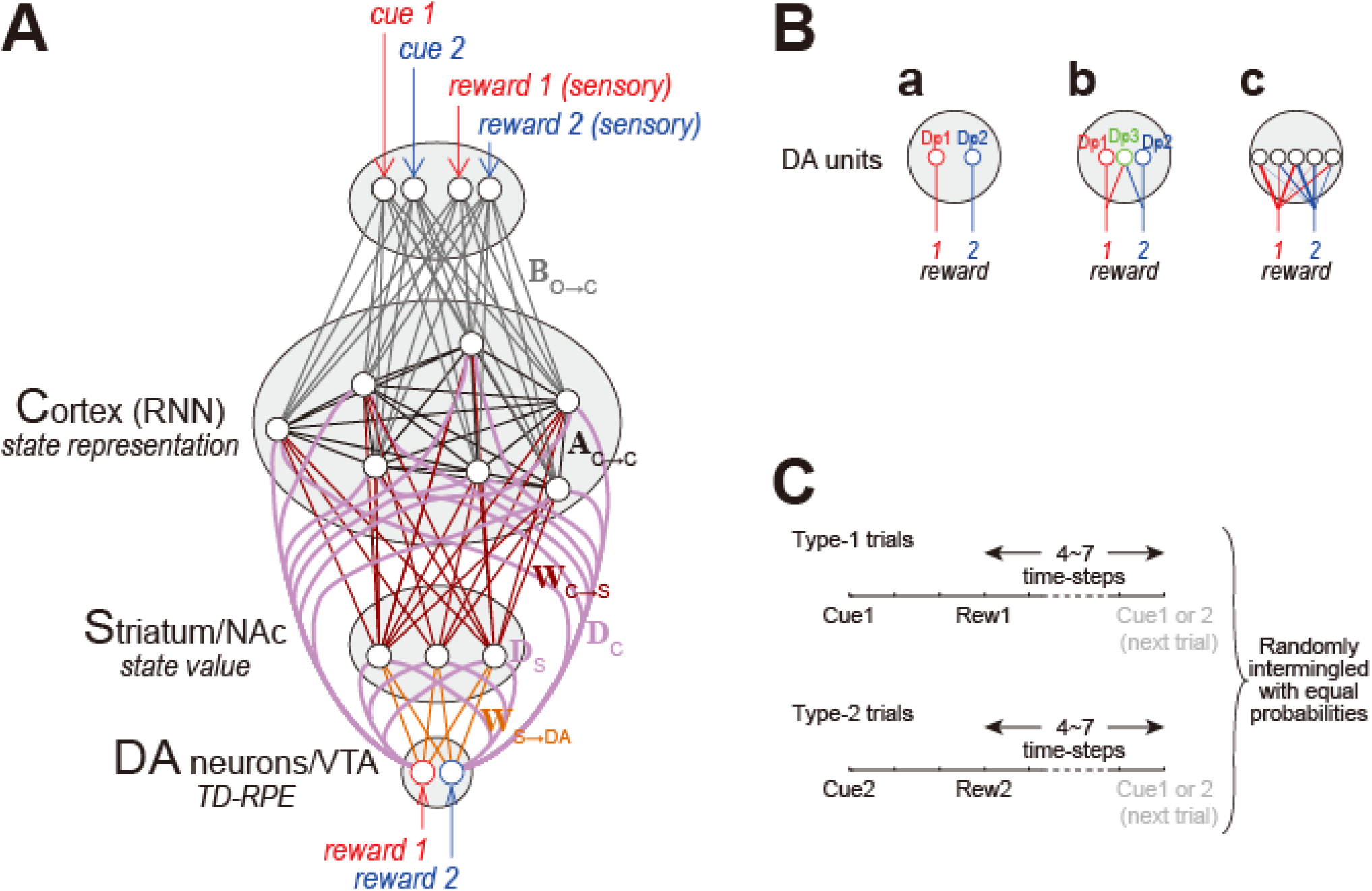
Schematic diagrams of the model and the task. **(A)** Schematic diagram of OVRNN-RB with a simple setting that there are two DA units, each of which receives input exclusively from one of two rewards. **(B)** Different settings about DA units. **(a)** The simple setting as in (A). **(b)** An extended setting, where an additional DA unit (Dp3) receives inputs evenly from both rewards. **(c)** The general setting, where multiple DA units receive inputs from the two rewards with fixed random weights. **(C)** Schematic diagram of the simulated behavioral task.

### Learning of reward-specific representation and value through double feedback alignments

We simulated a task with multiple cue-reward associations (Fig. 1C). There were two cues, Cue1 and Cue2, and two rewards, Rew1 and Rew2. There were two types of trials. In type-1 trials, Cue1 was presented, and three time-steps later, Rew1 was obtained, whereas in type-2 trials, Cue2 was presented, and three time-steps later, Rew2 was obtained (single time-step was assumed to correspond to several hundreds of milliseconds: see the Methods for details). Type-1 trials and type-2 trials were randomly intermingled with equal probabilities, and inter-trial intervals were randomly set to 4 ∼ 7 time-steps. We examined how the cortex(RNN)-striatum-DA connections and striatum-DA weights developed and the system behaved in the two types of trials.

We started with a simple configuration of the OVRNN-RB model where there were two DA units, Dp1 and Dp2, each of which received input exclusively from one of the two rewards, Rew1 and Rew2, respectively (Fig. 1Ba). To estimate the alignment of weights, we measured the correlation coefficient between the striatum-DA weights and the fixed random DA-striatum weights (denoted as *r*_SD&DS_, Fig. 2A) and the correlation coefficient between the cortex(RNN)-striatum-DA connections (i.e., product of the RNN-striatum weights and the striatum-DA weights) and the fixed random DA-RNN weights (denoted as *r*_CD&DC_, Fig. 2B). Across-simulation averages of *r*_SD&DS_ rapidly increased and then modestly decreased but remained positive, while *r*_CD&DC_ increased more gradually. These positive correlations indicate that alignments of the cortex-striatal-DA connections and the striatum-DA weights to the mesocortical and mesolimbic weights, respectively, both occurred.

**Figure 2.**
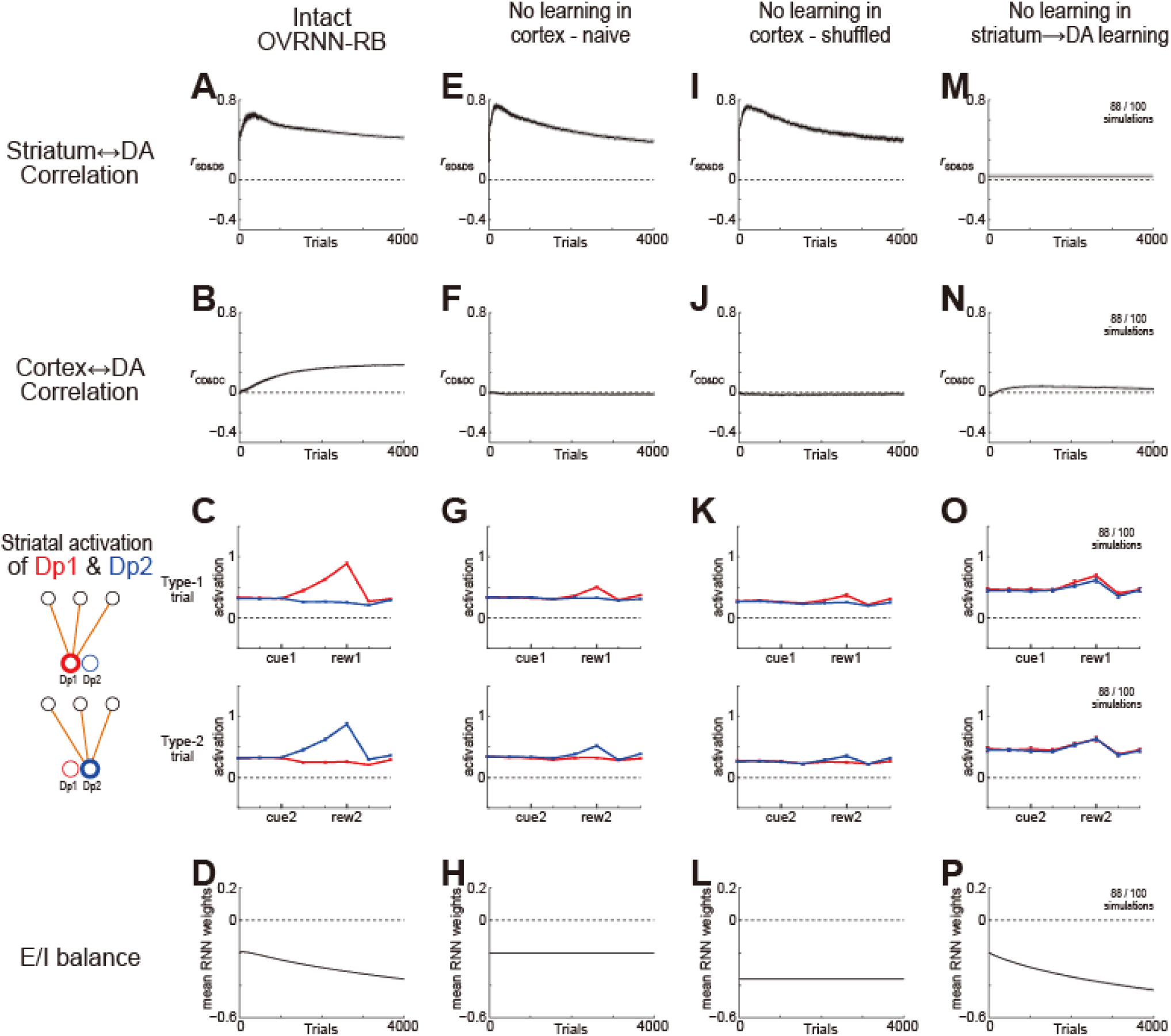
Behavior of OVRNN-RB with two DA units. **(A-D)** Results for the intact OVRNN-RB: (A,B) Across-trial evolution of the correlation coefficient between the striatum-DA weights and DA-striatum weights (*r*_SD&DS_) (A) or between the cortex(RNN)-striatum-DA connections (product of the cortex-striatum weights and the striatum-DA weights) and the DA-cortex(RNN) weights (*r*_CD&DC_) (B); (C) Striatal activations of the DA units (red: Dp1, blue: Dp2) (i.e., product of the striatum-DA weights and the striatal units’ activities) in the last type-1 trial (top) and type-2 trial (bottom) within 4000 trials; (D) Across-trial evolution of the E/I balance, i.e., the mean of the weights onto the RNN units. For all of (A-D), the averages over 100 simulations were shown as think black lines, and ±SEM were shown as thin gray lines (A,B,D) or error bars (C) (although almost invisible because SEM was small). **(E-H)** Results for a variant model with naive untrained-RNN, in which update of the weights onto the RNN units was omitted while all the other settings, including the initializations of the variables, were unchanged. **(I-L)** Results for another variant model with shuffled untrained-RNN, in which update of the weights onto the RNN units was omitted and the weights onto the RNN units were initialized to the values that were randomly shuffled from the weights in the leaned OVRNN-RB model at 4000-th trial. **(M-P)** Results for yet another variant model, in which learning of the striatum-DA weights was omitted and those weights were instead fixed to be random values. 12 out of 100 simulations in which the striatal activation of DA units became excessively high (>2) were omitted from plotting.

Figure 2C shows across-simulation averages of the striatal activations of DA (Dp1/Dp2) units (i.e., product of the striatum-DA weights and the striatal units’ activities) in the last type-1 trial (Fig. 2C-top) and type-2 trial (Fig. 2C-bottom) within 4000 trials, respectively. In type-1 trial, activation of Dp1 unit encoded the (temporally discounted) state values starting from Cue1 and ending upon Rew1 while activation of Dp2 unit was largely flat, whereas in type-2 trial, activation of Dp2 unit encoded the state values from Cue2 to Rew2 while activation of Dp1 unit was largely flat. These results indicate that reward-specific state representation and value were successfully learned in the model. Crucially, this enables reward-specific motivational control, as in the ancestor RB model ^36^. Specifically, under a situation where Rew1 (e.g., food) entails a high motivational desirability (i.e., hunger), the raised motivation for Rew1 can be encoded as a raised tonic DA from Dp1 unit, which enhances responses of striatal units that receive strong inputs from Dp1. Then, if these striatal units in return strongly project to Dp1, i.e., if the striatum-DA weights are aligned to the DA-striatum weights, the enhancement of their responses means a specific amplification of the values from Cue1 to Rew1 (i.e., values preceding food), without amplification of the values from Cue2 to Rew2.

To understand how the learning of the RNN weights contributed to these results, we examined two variant models: naive untrained RNN and shuffled untrained RNN. In both models, update of the RNN weights was omitted while update of the RNN-striatum and striatum-DA weights was kept intact. In the naive untrained RNN variant, RNN weights were initialized in the same manner as in the OVRNN-RB. In the shuffled untrained RNN variant, to initialize the RNN weights, we took the weights of a learned OVRNN-RB model (at 4000-th trial) and randomly shuffled those. Figures 2E-G and 2I-K show the results for the two variants. In both, *r*_SD&DS_ rapidly increased, indicating that alignment at the striatum-DA part still occurred, whereas *r*_CD&DC_ remained around 0, indicating no alignment (as expected). Trial-type-specific development of the activation of Dp1 or Dp2 unit was much poorer than the case of the intact OVRNN-RB model. These results indicate that alignment at the cortex(RNN)-striatum part requires learning of the RNN weights, and it is pivotal for development of reward-specific values when appropriate state representation is not given but needs to be learned in the RNN.

We also examined a third variant, in which update of the striatum-DA weights was omitted while update of the RNN and RNN-striatum weights was kept intact. In this case, in some simulations (12 out of 100), the striatal activation of DA units became excessively high (>2), indicating a learning failure. Analyzing the remaining simulations, *r*_SD&DS_ remained to be around 0 (Fig. 2M) as expected while *r*_CD&DC_ increased but only slightly (Fig. 2N), and state values were developed to a certain extent but there was no trial-type/unit-selectivity, also as expected (Fig. 2O).

Next, we examined an extended configuration of the OVRNN-RB model where there were three DA units (Fig. 1Bb): Dp1 and Dp2 receive exclusive inputs from reward Rew1 and Rew2, respectively, while Dp3 receives inputs evenly from both rewards. As shown in Fig. 3A,B, *r*_SD&DS_ rapidly increased and remained to be positive, and *r*_CD&DC_ gradually increased, similarly to the simpler case without Dp3 unit. As for the striatal activations of DA units, Dp1 and Dp2 units were activated in type-1 and type-2 trials, respectively, while Dp3 unit was activated to an intermediate level in both trial types (Fig. 3D). This indicates that reward-specific values were developed also in this model with three DA units. Finally, we examined a more general configuration where there were five DA units, which received fixed random (varied across simulations) inputs from the two reward types (Fig. 1Bc). In this model too, both *r*_SD&DS_ and *r*_CD&DC_ became positive (Fig. 3E,F), indicating the occurrence of double feedback alignments, although in 5 out of 100 simulations, the striatum-DA weights returned to **0** after 100 trials (i.e., learning failed and restarted), and they were omitted from the figures.

**Figure 3.**
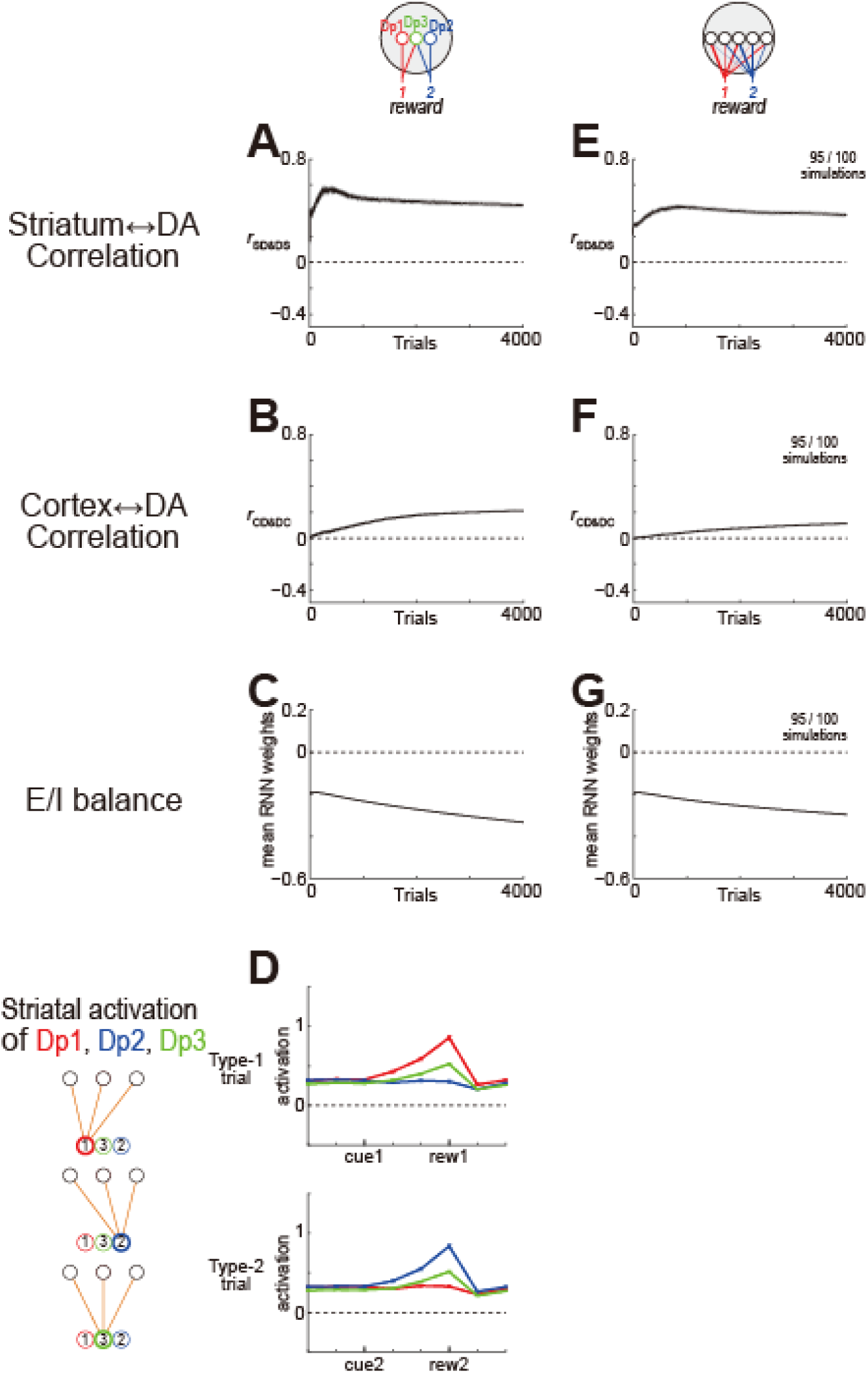
Behavior of OVRNN-RB with more than two DA units. **(A-D)** Case with three DA units: Dp1 and Dp2 receive inputs exclusively from Reward 1 and Reward 2, respectively, and Dp3 receives inputs evenly from both rewards. In (D), the green lines indicate the striatal activation of Dp3 unit. **(E-G)** Case with five DA units, which receive inputs from the two rewards with fixed random weights. The number on the top-right indicates the number of plotted simulations (out of 100 simulations) in which learning failure (weights returning to **0**) did not occur at least until 100-th trial (same applied to Fig. 4,7).

### Excessive excitation causes striatum-DA anti-alignment

In OVRNN-RB, the RNN weights were allowed to take both positive and negative values for simplicity. From the previous work, we can assume that the model also works in a more biologically plausible setting where output weights of each unit can only be either excitatory and inhibitory ^41^. In all the models, the mean RNN weight started from a negative value because of the inhibition-dominant initialization (mean weight: −0.2), and training made the RNN weights further negative (Fig. 2D,P and Fig. 3C, G) (although initially a slight positive shift appeared), unless the update of the RNN weights was omitted (Fig. 2H,L).

To test how important it was to initialize the RNN with negative average weights, we initialized the RNN weights to be excitation-dominated (i.e., the initial mean RNN weight to be 0.1 instead of −0.2 that was so far assumed). With such excitation-dominated RNNs, there were many occasions (51 out of 100 simulations) when learning failed (i.e., the striatum-DA weights returned to **0**) after 100 trials. In the remaining cases where the model successfully learned, *r*_SD&DS_ rapidly decreased to become negative, and then increased to eventually become positive (Fig. 4A), while *r*_CD&DC_ slowly increased (Fig. 4B). The initial decrease and negative value of *r*_SD&DS_ were also observed in the model with native untrained RNN with excitation-dominant initialization (Fig. 4D). In contrast, such a pattern did not appear (Fig. 4G) in the model with untrained RNN, whose weights were shuffled from learned OVRNN-RB, which was initialized to be excitation-dominant but eventually became inhibition-dominant (as shown in Fig. 4C).

**Figure 4.**
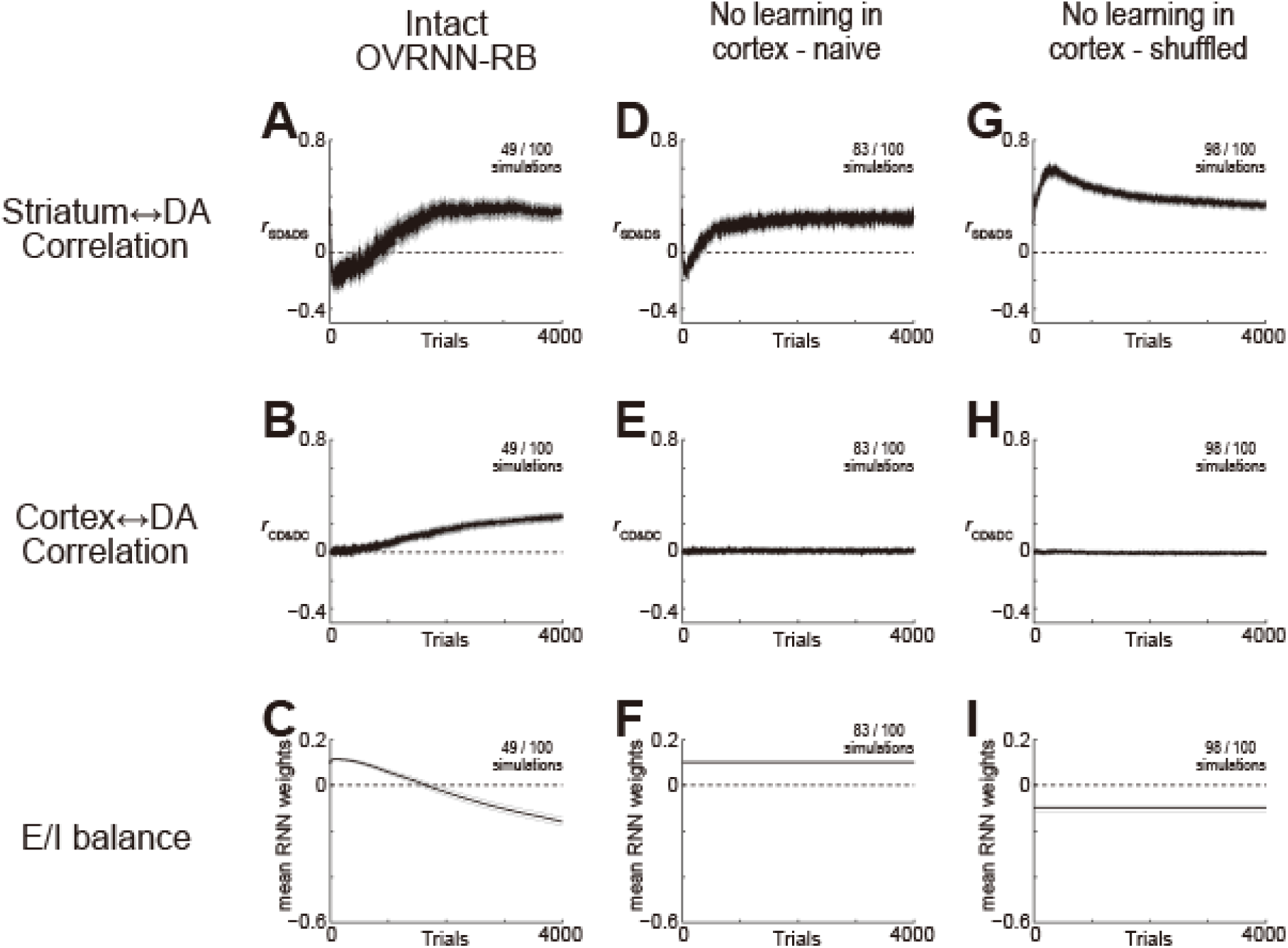
Behavior of OVRNN-RB with two DA units, with the E/I balance was initialized to be excitation-dominant. **(A-F)** Results for the intact OVRNN-RB (A-C) or the model with naive untrained RNN (D-F) with the mean weight onto the RNN units was initialized to 0.1 (instead of −0.2 as done in Fig. 2,3). **(G-I)** The model with shuffled untrained RNN, whose weights were shuffled from the leaned intact OVRNN-RB with excitation-dominant initialization (i.e., those shown in (A-C)) and fixed.

These results suggest that negative *r*_SD&DS_, i.e., anti-alignment of the striatum-DA weights to the DA-striatum weights may be caused by E/I imbalance in the cortical RNN, more specifically, excessive excitation (or insufficient inhibition). Such an anti-alignment should impair reward-specific motivational control, because it would mean a decrease in the value of states associated with highly desired rewards while state values leading to other rewards are amplified (e.g., when hungry, states leading to food are lower valued whereas values leading to drink are higher valued, and opposite occurs when thirsty).

### Role of persistent activity

To better understand the possible anti-alignment in excitation-dominated networks, we conducted simulations using OVRNN-RB with a simplified setting, where two DA units (Dp1 and Dp2), activated exclusively by Rew1 and Rew2, project exclusively to two striatal units (St1 and St2), respectively (Fig. 5A), with the RNN initialized to either inhibition-dominant or excitation-dominant. In the case of inhibition-dominant initialization, St1-Dp1 and St2-Dp2 weights became stronger than St1-Dp2 and St2-Dp1 weights (i.e., the striatum-DA weights were aligned to the fixed DA-striatum weights) within 200 trials (Fig. 5B). In contrast, in the case of excitation-dominant initialization, St1-Dp2 and St2-Dp1 weights became stronger than St1-Dp1 and St2-Dp2 weights, i.e., the striatum-DA weights were anti-aligned to the fixed DA-striatum weights (Fig. 5C). Positive mean value of weights can affect the activity of RNNs in multiple ways, e.g., the dynamics can become unstable or network reaches a trivial fixed point with persistent activity ^48^. We hypothesized that emergence of persistent activity in excitation-dominated networks may underlie the anti-alignment.

**Figure 5.**
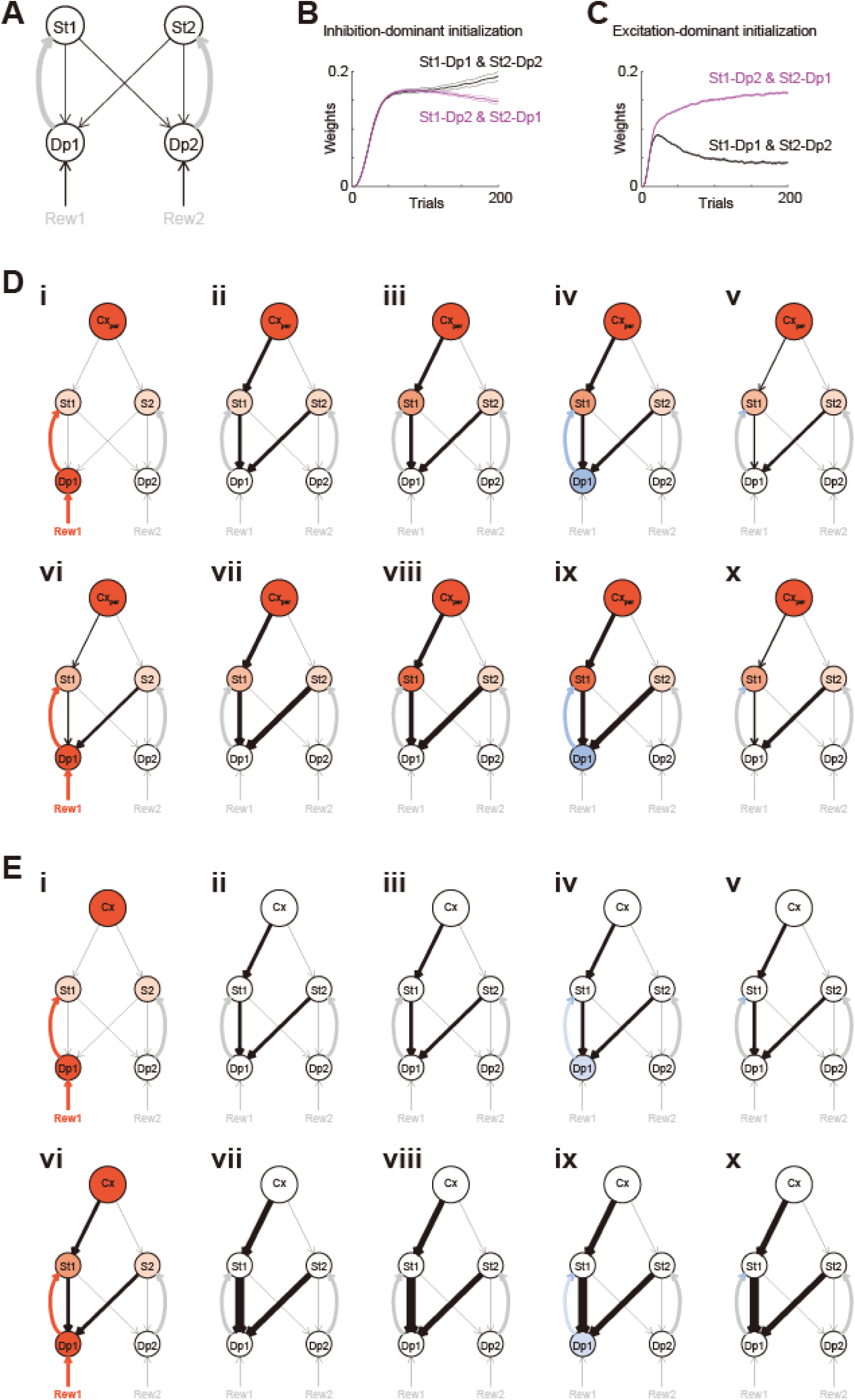
Excessive excitation and persistent activity in the cortex (RNN) causes anti-alignment of the striatum-DA weights to the DA-striatum weights. **(A)** OVRNN-RB with a simple setting, where two DA units (Dp1 and Dp2), activated exclusively by Rew1 and Rew2, project exclusively to two striatal units (St1 and St2), respectively. **(B,C)** Across-trial evolution of the mean strength of St1-Dp1 & St2-Dp2 (black) and St1-Dp2 & St2-Dp1 (magenta) connections in the case of inhibition-dominant (−0.2) (B) or excitation-dominant (0.1) (C) initialization of the RNN weights. **(D,E)** Schematic explanation of how cortical persistent activity (Cx_per_) causes anti-alignment (i.e., St1-Dp1 < St1-Dp2) (D) whereas how positive alignment (i.e., St1-Dp1 > St1-Dp2) is formed if there is no persistent cortical activity (E).

To illustrate this hypothesis, we used a simple configuration of our model with a persistently active cortical population (Cx_per_), two DA units (Dp1 and Dp2), and two striatal units (St1 and St2) (Fig. 5Di). In this model, we can easily understand how learning would shape the striatum-DA connections (i.e., St1→Dp1, St1→Dp2, St2→Dp1, and St2→Dp2) if they all had the same initial weights.

When reward Rew1 is obtained, Dp1 encodes positive TD-RPE (Fig. 5Di), which causes the same level of potentiation of St1→Dp1 and St2→Dp1 weights, and also potentiates Cx_per_→St1 but not Cx_per_→St2 weight via the exclusive Dp1→St1 connection (Fig. 5ii): thus, St1 becomes more activated by Cx_per_ than St2 (Fig. 5Diii). In the subsequent time-steps without reward, Dp1 tends to encode negative TD-RPE (Fig. 5Div), because TD-RPE = 0 + *γ*·value(*t*) − value(*t*−1) tends to be negative given that the time discount factor *γ* is smaller than 1 and the value function has not been well learned so that value(*t*) and value(*t*−1) take similar random values. This negative TD-RPE causes depression (LTD, or depotentiation) of St1→Dp1 and St2→Dp1 weights, but crucially, more prominently for St1→Dp1 than St2→Dp1 weights (Fig. 5Dv) because St1 is more activated by Cx_per_ than St2. As a result, St1→Dp1 weight becomes weaker than St2→Dp1 weight (Fig. 5Dv).

At the next occasion of reward Rew1 receival, Dp1’s positive TD-RPE (Fig. 5Dvi) causes potentiation (LTP) of St1→Dp1 and St2→Dp1 weights, this time more prominently for St1→Dp1 than St2→Dp1 weights (differently from the initial occasion shown above) because St1 is more active than St2 (Fig. 5Dvii), mildening the previously formed difference between St1→Dp1 and St2→Dp1 weights. However, this positive TD-RPE further potentiates Cx_per_→St1 but not Cx_per_→St2 weight, and in the subsequent time-steps without reward, St1 is even more activated by Cx_per_ than St2 (Fig. 5Dviii) so that negative TD-RPE causes depression (LTD, or depotentiation) of St1→Dp1 and St2→Dp1 weights, even more prominently for St1→Dp1 than St2→Dp1 weight (Fig. 5Dix). Therefore, in total, St1→Dp1 weight would become even weaker than St2→Dp1 weight (Fig. 5Dx). Repeating these processes, also similarly for reward Rew2 and Dp2 unit, St1→Dp1 and St2→Dp2 weights would become weaker than St1→Dp2 and St2→Dp1 weights, explaining the formation of anti-alignment.

In contrast, if there is no persistent cortical activity after reward, upon the first receival of reward Rew1 (Fig. 5Ei-v), positive TD-RPE potentiates St1→Dp1 and St2→Dp1 weights equally, and also potentiates Cx_per_→St1 weight. Then, upon the second receival of reward Rew1 (Fig. 5Evi-x), positive TD-RPE potentiates St1→Dp1 more than St2→Dp1 weights because St1 is more active than St2. In this way, positive alignment will be formed.

### Excitation/inhibition imbalance induced by different processes causes similar learning impairment

In the simulation shown in Fig. 4A-C where the RNN weights were initialized to be excitation-dominant, more specifically, 0.1 on average, even though anti-alignment of the striatum-DA and DA-striatum weights (i.e., negative *r*_SD&DS_) occurred in the early phase, *r*_SD&DS_, as well as *r*_CD&DC_, eventually became positive, indicating successful learning, and the mean RNN weights eventually became negative. We examined learning in the case where the RNN weights were initialized to be more excitation-dominant, in particular, 0.2 on average. As shown in Fig. 6A-C, in this case, *r*_SD&DS_ remained to be negative and *r*_CD&DC_ remained to be around 0 for a long duration, indicating the persistence of learning impairment, although they still continued to increase slowly while the mean RNN weights continued to decrease.

**Figure 6.**
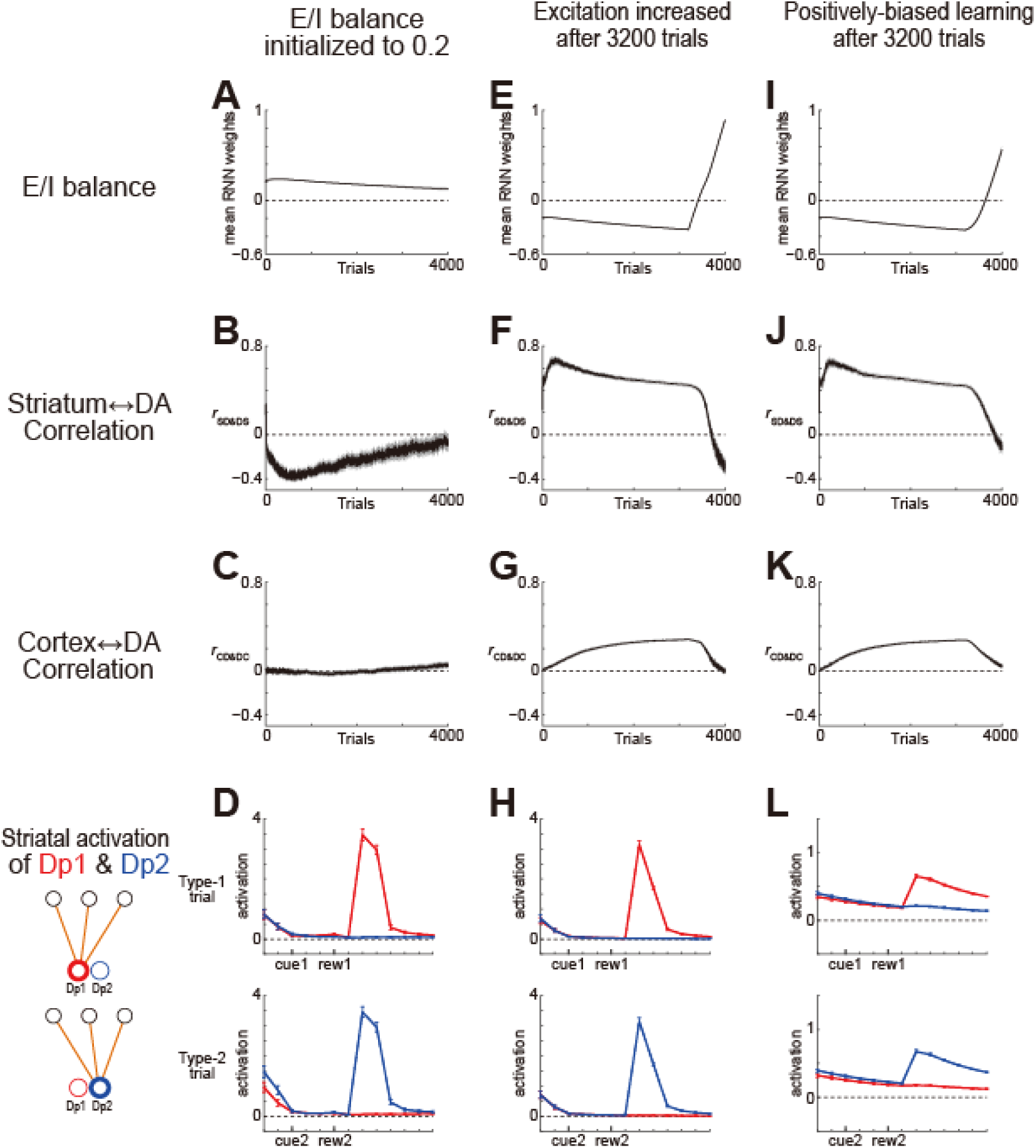
Effects of manipulations that affect the E/I balance. **(A-D)** The E/I balance was initialized to be more excitation-dominant (mean RNN weight = 0.2). **(E-H)** After 3200 trials, a small positive value (0.0002) was added to each RNN weight at each time-step. **(I-L)** After 3200 trials, the learning rate for the RNN weights was doubled when DA received at post-synaptic unit (i.e., product of RPE and the DA-striatum connections) was positive and halved when it was negative. In this Fig. 6, at each trial, simulations where correlation coefficient could not be calculated (because all the elements of either weight vector had a same value such as 0) were omitted from plotting, while simulations with learning failure within 100 trials were included (different from Fig. 3,4,7).

Next, in order to model developmental and/or environmental factors that affect the E/I balance, we examined the cases where the RNN weights were initialized to be inhibition-dominant and learning proceeded properly but later on manipulation was added so that the RNN weights were shifted to the positive direction. We examined two sorts of manipulations. The first one was a direct positive shift of each RNN weight, which could abstractly model glutamate or GABA-related factors. Specifically, a small positive value (0.0002) was added to each RNN weight at each time-step. The second manipulation was a bias in the learning rate for the update of the RNN weights, which was based on the suggestions that changes in the DA level could cause learning rate biases, specifically, larger learning rate for positive than negative, or negative than positive, RPE ^37, 49, 50^, although such suggestions were made about corticostriatal weights rather than about intra-cortical weights. Specifically, we considered a manipulation where the learning rate was doubled when DA received at post-synaptic unit (i.e., product of RPE and the DA-striatum connections) was positive and halved when it was negative.

Figure 6E-G and 6I-K show the results of simulations, in which either of the two manipulations was applied after 3200 trials. Both manipulations shifted the mean RNN weights to the positive direction, as expected (Fig. 6E,I). The correlation between the striatum-DA and DA-striatum weights (*r*_SD&DS_) decreased and eventually became negative (Fig. 6F,J), indicating that their alignment, necessary for reward-specific motivational control, was degraded and eventually reversed. The correlation between the cortex(RNN)-striatum-DA connections and DA-cortex(RNN) weights (*r*_CD&DC_) also decreased and approached around 0 (Fig. 6G,K), indicating that their alignment was also degraded. These results demonstrated that even after learning of OVRNN-RB properly proceeded and both alignments were formed, the alignments were degraded by either a direct positive shift of the RNN weights or a positive bias in the learning rate for their update.

Figures 6D,H,L show the striatal activations of the DA units in the second last type-1 and type-2 trials within 4000 trials in the cases where the RNN weights were initialized to excitation-dominant (0.2) (Fig. 6D) or either of the two manipulations was applied after 3200 trials (Fig. 6H,L). Compared to the results without manipulation (Fig. 2C) where there appeared activation spanning from a cue to a reward, which means that the credit of the reward was assigned to the preceding states after the cue, no such activation was formed and instead activation following reward receival was formed, indicating that correct credit-assignment was failed and spurious credit assignment was formed.

## Discussion

In order to mechanistically understand the roles of mesocorticolimbic DA in value learning, motivational control, and cognitive functions, we developed a neurocomputational model, combining previous models that explained either reward-specific motivational control or learning of context-dependent state representation and value but not them together. Our combined model, OVRNN-RB, provides an integrated account of all these functions. A key factor for the model’s proper operation turned out to be the E/I balance in the cortical RNN. Change in the balance toward excessive excitation, whether innately existing or caused by different manipulations, degrades or even reverses the alignments of cortical and striatal downstream connections to the DA feedback weights, causing impairments in reward-specific motivational control and credit assignment. As we discuss below, this provides an integrated account for diverse symptoms and suggested causes of schizophrenia.

### Normal operation of OVRNN-RB through double alignments

Normal operation of OVRNN-RB is realized through the occurrence of double alignments. Reward-specific motivational control is achieved through the alignment of the learnable striatum-DA weights to the randomly fixed DA-striatum (mesolimbic) feedback weights, as schematically explained in the Results (in the explanation of Fig. 2C), as in the ancestor RB model ^36^. Meanwhile, reward-specific credit assignment is achieved through the alignment of the learnable cortex(RNN)-striatum-DA connections to the randomly fixed DA-cortex(RNN) (mesocortical) feedback weights by virtue of the TD-learning’s capability of solving credit assignment, as in the ancestor OVRNN model^41^. Demonstration of the occurrence of such double feedback alignments in RL of RNN and its downstream trained by online TD-RPE is a novel achievement of the present work, although occurrence of feedback alignments at multiple stages in supervised learning of the feed-forward network was previously shown ^44^.

### Negative symptoms of schizophrenia explained by degraded/reversed striatum-DA alignment

As explained in the Results, the alignment of striatum-DA forward and feedback weights enables reward-specific motivational control ^36^: when a particular reward, e.g., food, has a high motivational value, i.e., in hunger, responses of striatal units that are strongly innervated by DA units activated by food become enhanced by tonic DA, and if these striatal units in return strongly project to the food-activated DA units (i.e., alignment occurs) and thereby encode state/action values leading to food, the enhancement of their responses means an enhancement of the state/action values leading to food, which promotes food intake.

If the striatum-DA alignment is degraded or even reversed, even though food should have a high motivational value, i.e., objectively in hunger, the state/action values leading to food may not be enhanced and can even be diminished, and so food intake may not be promoted but even decreased. This could manifest as a lack of motivation (avolition) or anhedonia, which are major constructs of negative symptoms ^21, 22^. In the meantime, the opposite can also occur. Specifically, even though food should not have a high motivational value, i.e., objectively in satiety, the state/action values leading to food may not be diminished and can even be enhanced, and so food intake may not be decreased but even promoted. This would cause overeating and could lead to obesity, potentially explaining that problems in food intake, including obesity, are frequently seen in schizophrenia patients ^51^.

### Positive symptoms of schizophrenia explained by impaired credit assignment

The positive symptoms of schizophrenia, delusion and hallucination, could originate from problems in credit assignment (c.f., ^20^), i.e., difficulties in assigning appropriate credits for a particular event to actions/states that caused or lead to the event. Since OVRNN-RB achieves credit assignment, impairment of OVRNN-RB could potentially explain the positive symptoms. However, while OVRNN-RB achieves credit assignment for reward values, what typically matters in delusion or hallucination is credit assignment for actions or intentions. But this gap can be filled by a recent finding of distinct DA activity that encodes action prediction error (APE) instead of RPE ^52^. If DA units in OVRNN-RB encode APE instead of RPE, credit assignment for actions, or intentions as internal (non-motor) actions, could potentially be achieved.

APE-encoding DA signals were found in a particular part of the striatum, namely, the tail of striatum (TS) of mice engaging in an auditory discrimination task ^52^. TS receives inputs from auditory, visual, and other many cortical and subcortical regions ^53^, but in a previous study ^54^, among cortical areas that were examined, the proportion of TS-projecting neurons was highest in the auditory cortex (∼25%) and lower in the visual cortex (∼12∼3%) (Figure 2F of ^54^), and TS was also shown to relay auditory signals ^55^. It may be of importance to explore if APE-encoding DA signals in TS are particularly associated with auditory stimuli or similarly associated with visual and other sensory stimuli, given that auditory hallucination is the most common in schizophrenia ^56^.

Other than APE-encoding DA signals, TS is also known to host a different kind of non-RPE-encoding DA signals, specifically, those encoding prediction errors of potential threat or salience ^57-60^. Given this, it may be valuable to explore if problems in credit assignment of potential threat could account for certain aspects of symptoms. More generally, there appears to be a possibility that diverse symptoms and impairments in schizophrenia could in part be understood by which striatal/cortical regions and DA populations targeting there are affected.

### Dopamine, glutamate, and developmental causes: different origins converging onto E/I imbalance

At the pathophysiological level, there are three prominent hypotheses that explain majority of schizophrenia ^31, 32^. The DA hypothesis is based on the psychomimetic effects of DA receptor agonists ^18^ and antipsychotic effects of antagonists ^19^. The glutamate hypothesis is based on the psychomimetic effects of NMDA receptor antagonists ^61^, which could result in pyramidal neuronal firing ^62^ via regulation of inhibitory interneurons. The developmental hypothesis ^63-65^ is based on changes in synapse formation and elimination by sociodevelopmental factors. These hypotheses have been proposed to be potentially integrated into changes in the E/I balance ^32-35^.

However, while previous modeling studies revealed how the E/I imbalance impairs cortical functions ^66-71^, its interaction with RPE-based RL in DA-basal ganglia circuits remained to be explored. Usually, functional role of E/I balance is understood from a ‘bottom-up’ viewpoint, i.e., in the way it changes the network dynamics (e.g. oscillations, synchrony) and the gain of neurons. Here we take a complementary ‘top-down’ (task-variable-based c.f. ^72^) perspective on the functional role of E/I balance. We have shown that in our OVRNN-RB, initial or introduced excessive excitation similarly disrupted the functions (formation of state values (task variables)) by creating misalignment of synaptic connectivity crucial for learning. In our model, we changed E/I balance by changing the mean of the RNN weights. This kind of E/I balance is referred to as loose global balance ^73^. This is of course simplistic as E/I balance in schizophrenia could have a more complicated dynamics, as suggested in the previous studies ^66-71^. Yet, a change in loose global balance could correspond to changes in the synaptic connectivity due to genetic and sociodevelopmental factors. We have also shown that introduction of positive bias in the learning rate also lead to similar dysfunctions, and this could potentially model the effect of excessive DA.

Regarding how the E/I imbalance causes symptoms, it was proposed ^32^ that increased "noise" could cause functional impairments, but exact mechanisms remain elusive. Our model describes how the E/I imbalance impairs reward-specific motivational control and credit assignment through degradation of the alignments, providing a concrete mechanistic explanation.

### RL-based understanding of schizophrenia

Previous studies linked multiple aspects of schizophrenia to RL impairments and their neural substrates ^24-26, 74^: (i) flexible goal-directed RL was impaired whereas simple RL in stable environments was spared ^24, 27^, (ii) impaired performance in RL tasks can in fact come from working memory deficits ^28-30^, (iii) representation of value ^24^ or state ^75^ may be impaired, and (iv) the cortico-striatal circuits, in particular, the ventral striatum (VS) and the prefrontal/orbitofrontal cortex (PFC/OFC) may be affected ^26^. These findings are broadly in line with our model. Specifically, OVRNN, modeling the cortico-striatal system with cortical DA projections such as PFC/OFC-VS, learns adaptive state representation and value, while value learning itself is possible even if state representation is not learned but given by neural representation of sensory inputs in simple stable situations. Also, the hypothesized cortical excessive excitation and persistent activity impaired the formation of state representation bridging the cue and reward, which is a sort of working memory, and would also impair working memory functions more generally.

The previous RL-based studies using model fitting and comparison have precisely characterized certain aspects of schizophrenia. Complementary to these, our model tries to explain the whole picture of schizophrenia in the RL framework, integrating the diverse symptoms and causes. It was enabled by incorporation of reward identity/diversity (RB) and state representation learning (OVRNN), which were missing in conventional RL models (and tasks). Even though different from the transdiagnostic dimensional approach ^76^, first-principle models could greatly help elucidation of the mechanisms of disorders, as exemplified for addiction ^77-79^.

### Predictions

Here we present two lines of predictions of our model. First, the defining feature of OVRNN-RB is the occurrence of double alignments of the striatum-DA and cortex-striatum-DA pathways to the mesolimbic and mesocortical DA weights, respectively. The striatum-DA alignment precedes (compare Fig. 2A and 2B), and it might be completed during developmental stages. The cortex-DA alignment occurs gradually, presumably during learning of behavioral tasks. Most simply, OVRNN-RB predicts that if a local cortical cite receives strong DA input upon a particular reward over other rewards, the forward connection from that cite (via striatum) to DA neurons that show strong response to that specific reward, measured as the DA neurons’ response to a simulation of that cortical cite, gradually increases in the course of learning. Moreover, such an increase should be impaired when the cortical cite is manipulated to entail persistent activity, or schizophrenia model animals, for which such persistent activity is expected to be observed. These could be tested in the prefrontal cortex, but also in the hippocampus, where disruption of reward representation in rat model of schizophrenia risk was experimentally suggested ^80^.

Second, the key suggestion of our model is that cortical persistent activity due to excessive excitation degrades/reverses the alignment of striatum-DA connections, resulting in functional impairments. Because update of the RNN weights was assumed to depend on DA-encoding TD-RPE: **C**_RD_***r***(*t*) + *γ***W**_SD_***v***(*t*+1) − **W**_SD_***v***(*t*) (where *γ* is the time discount factor; see the Methods for details), if *γ* is smaller, i.e., temporal discounting is severer, TD-RPE, and thereby the RNN weight update would tend to be more negative, preventing persistent activity. Indeed, when *γ* was changed from the original 0.8 to 0.7 in the simulation with the mean RNN weight initialized to 0.1, simulations with a learning failure in the initial 100 trials became less frequent (51 to 31 out of 100), and in the remaining simulations, negative correlation (anti-alignment) between the striatum-DA and DA-striatum weights turned into positive at earlier timings (compare Fig. 7A and 7D). As such, our model predicts that the degree of temporal discounting is *negatively* associated with cortical excessive excitation and functional impairments. Notably, while temporal discounting is *positively* associated with several other psychiatric disorders, results for schizophrenia were mixed ^81^ with either positive ^82^ or negative ^83^ association reported. The latter result, though potentially due to technical issues discussed in that study, is in line with our prediction, and further tests, including those using shorter delays (on the order of seconds) and examining/manipulating the E/I balance in animal models, are desired to be executed.

**Figure 7.**
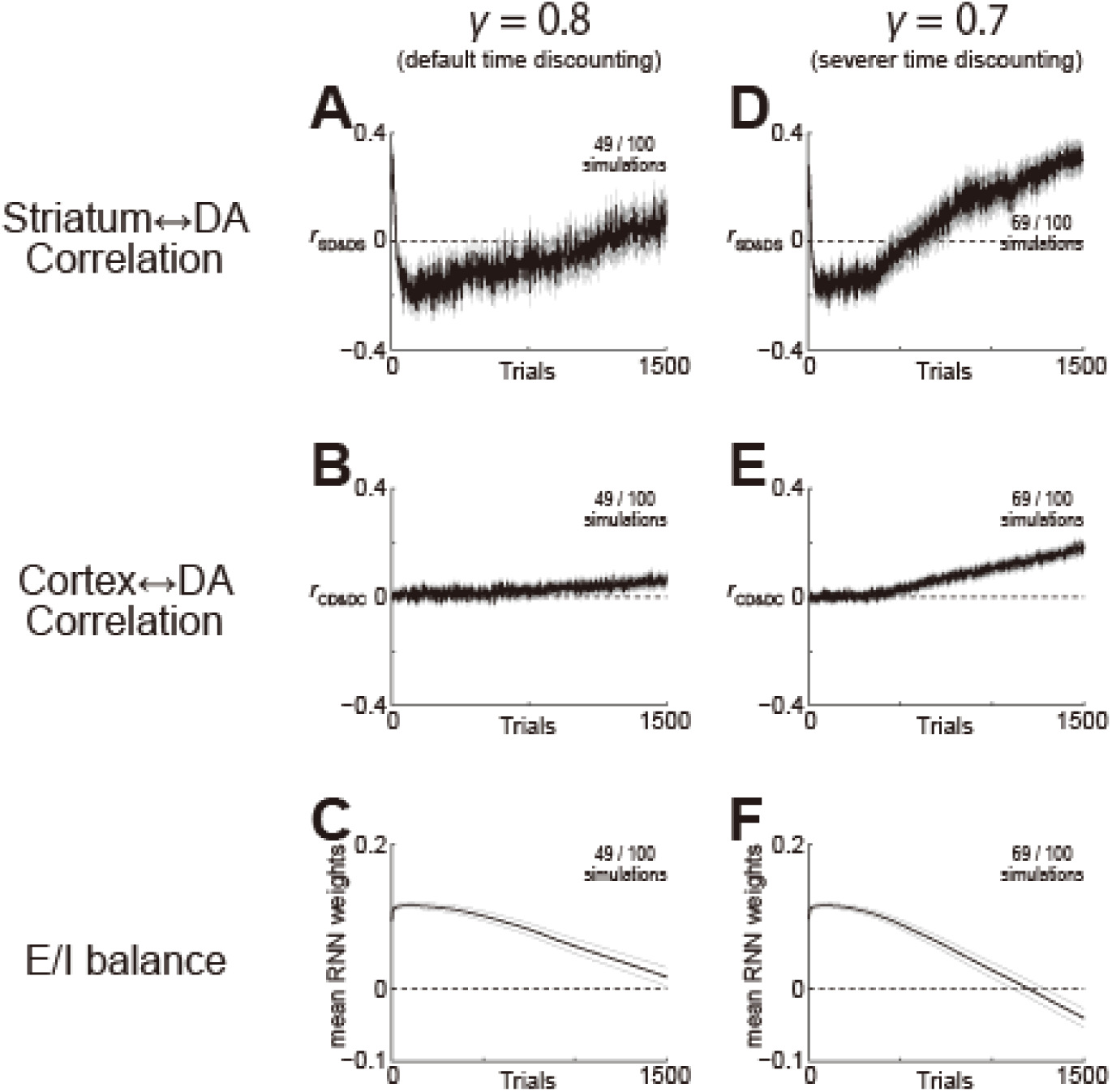
Effects of temporal discounting on the dynamics of OVRNN-RB with two DA units, with the E/I balance initialized to be excitation-dominant (mean RNN weight = 0.1). **(A-C)** Results with the default value of time discount factor (*γ* = 0.8): enlarged views of 1st∼1500th trials of the results shown in Fig. 4A-C. **(D-F)** Results with a smaller time discount factor (*γ* = 0.7) (i.e., severer temporal discounting).

## Methods

### Online Value-RNN with Reward Bases (OVRNN-RB)

We constructed OVRNN-RB by combining OVRNN, specifically its version with random feedback and biological constraints ("oVRNNrf-bio" in ^41^), and RB ^36^. OVRNN-RB consists of the observation units (modeling sensory cortex), an RNN (modeling prefrontal/association cortex), striatal units, and DA units (Fig. 1A). As in OVRNN ^41^, each unit was assumed to represent a population of neurons, and single time-step was assumed to correspond to several hundreds of milliseconds, which are similar to the time scales of certain types of short-term synaptic plasticity ^84-86^ and the behavioral time-scale synaptic plasticity ^87, 88^ (see ^41^ for detailed discussion for these).

The activities of the observation units at time-step *t*, ***o***(*t*) = (*o_h_*(*t*)) (*h* = 1, …, 4), encoded the presence of a cue or a reward. Specifically, *o*_1_(*t*) or *o*_2_(*t*) became 1 when a cue, Cue 1 or Cue 2, was presented, respectively, while *o*_3_(*t*) or *o*_4_(*t*) became 1 when a reward, Rew 1 or Rew 2, was obtained, respectively, and at other time-steps *o_h_*(*t*) was set to 0.

The activities of the RNN units, ***x***(*t*) = (*x_j_*(*t*)) (*j* = 1, .., 40), were determined depending on the activities of themselves and the observation units at the previous time-step:

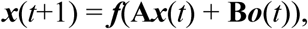

where **A** = (*A_ij_*) was the strength of the recurrent connection from *x_j_* to *x_i_* and **B** = (*B_ih_*) was the strength of the feed-forward connection from *o_h_* to *x_i_*. *f*(*z*) = 1/(1 + exp(−*z*)) was a sigmoidal function that represented the neuronal input-output relation.

The activity of the striatal units, ***v***(*t*) = (*v_k_*(*t*)) (*k* = 1, 2 (for Fig. 5) or 1, …, 10 (otherwise)), were determined by:

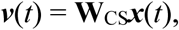

where **W**_CS_ = (*W*^CS^_*ki*_) was the cortex(RNN)-striatum weight from *x_i_* to *v_k_*.

The activities of the DA units, ***d***(*t*) = (*d_ℓ_*(*t*)) (*ℓ* = 1, 2 (for Fig. 2, 4-7) or 1, 2, 3 (for Fig. 3A-D) or 1, …, 5 (for Fig. 3E-G)), were determined by:

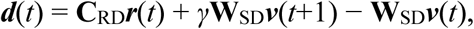

where ***r***(*t*) = (*r_m_*(*t*)) (*m* = 1, 2) encoded two rewards: *r*_1_(*t*) or *r*_2_(*t*) became 1 when Rew 1 or Rew 2 was obtained, respectively, and otherwise *r_m_*(*t*) = 0. **C**_RD_ = (*C*^RD^*_ℓm_*) was the fixed reward-DA weight from *r_m_* to *d_ℓ_*, and was set to

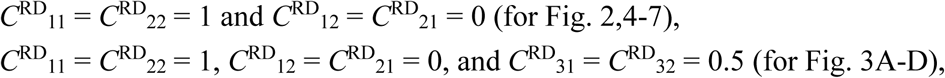

or each *C*^RD^*_ℓm_* was set to a pseudo uniform random number on [0 1] (for Fig. 3E-G). **W**_SD_ = (*W*^SD^*_ℓk_*) was the striatum-DA weight from *v_k_* to *d_ℓ_*, and *γ* was the time discount factor and was set to 0.8 except for the simulations shown in Fig. 7D-F, for which *γ* was set to 0.7. As such, ***d***(*t*) encoded the two-dimensional TD-RPE.

### Update rules of OVRNN-RB

The striatum-DA weight **W**_SD_ = (*W*^SD^*_ℓk_*) was initialized to 0 and updated according to:

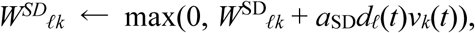

where max(*z*_1_, *z*_2_) returned the larger one of *z*_1_ and *z*_2_ (i.e., *W^SD^_ℓk_* was constrained to be non-negative) and *a*_SD_ was the learning rate and was set to 0.03.

The cortex(RNN)-striatum weight **W**_CS_ = (*W*^CS^_*ki*_) was also initialized to 0 and updated according to:

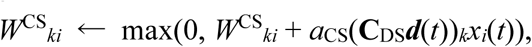

where the max operation ensured that *W*^CS^*_ki_* was also constrained to be non-negative. **C**_DS_ = (*C*^DS^_*kℓ*_) was the fixed DA-striatum weight, and (**C**_DS_***d***(*t*))*_k_* indicates the *k*-th element of **C**_DS_***d***(*t*). *a*_CS_ was the learning rate and was set to 0.03.

The recurrent and feed-forward connection strengths **A** = (*A_ij_*) and **B** = (*B_ih_*) were initialized to pseudo standard normal random numbers plus an offset (initial mean RNN weight), which was set to −0.2 (inhibition-dominant initialization: Fig. 2,3,5B,6E-L), 0.1 (excitation-dominant initialization: Fig. 4,5C,7), or 0.2 (more excitation-dominant initialization: Fig. 6A-D), and updated at every time-step as:

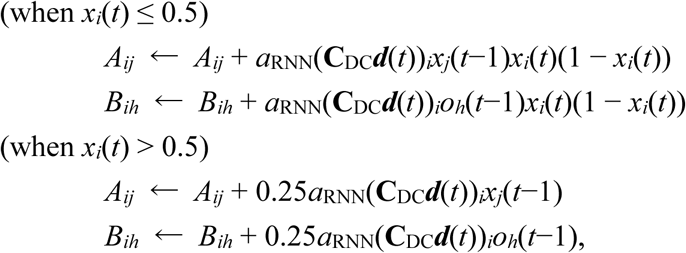

where **C**_DC_ = (*C*^DC^*_iℓ_*) was the fixed DA-cortex(RNN) weight, and (**C**_DC_***d***(*t*))*_i_* indicates the *i*-th element of **C**_DC_***d***(*t*). *a*_RNN_ was the learning rate and was set to 0.1 except for the time-steps after 3200 trials in Fig. 6I-L, for which *a*_RNN_ was set to 0.2 for *i* with the *i*-th element of **C**_DC_***d***(*t*) ≥ 0 and 0.05 for other *i* (i.e., doubled when DA received by the post-synaptic unit was positive and halved when it was negative (manipulation of learning rate bias)). The dependence on *x_i_*(*t*) (i.e., post-synaptic activity) was taken from the OVRNN model ^41^, where it was modified from the original non-monotonic dependence derived from gradient-descent so as to be monotonic + saturation. For the time-steps after 3200 trials in Fig. 6E-H, 0.0002 was added to each of *A_ij_* and *B_ih_* at every time-step (manipulation of excitation increase).

Each element of the fixed DA-striatum weight **C**_DS_ = (*C*^DS^*_kℓ_*) was set to a pseudo uniform random number on [0 1], except for the cases shown in Fig. 5 where *C*^DS^_11_ = *C*^DS^_22_ = 0.25 and *C*^DS^_12_ = *C*^DS^_21_ = 0. Each element of the fixed DA-cortex(RNN) weight **C** = (*C*^DC^_*kℓ*_) was set to a pseudo uniform random number on [0 1].

### Simulation of the behavioral task

We simulated a task of multiple cue-reward associations (Fig. 1C), with two cues, Cue1 and Cue2, and two rewards, Rew1 and Rew2. There were two types of trials, which were randomly intermingled with equal probabilities. In type-1 trials, Cue1 was presented and three time-steps later Rew1 was obtained, whereas in type-2 trials, Cue2 was presented and three time-steps later Rew2 was obtained. The cue or reward time-step marked in the figures was defined to be the time-step when the RNN received the cue or reward observation, respectively: if *o_h_*(*t*) = 1 at time *t*, *t* + 1 was defined to be a cue or reward time-step, respectively. Inter-trial interval (ITI) was set to 4, 5, 6, or 7 time-steps with equal probabilities. For each model and each condition, we conducted 100 simulations (with different pseudo-random numbers) of 4000 trials.

### Analyses, software, and code availability

Alignments of the forward connections to the feedback connections were evaluated by their correlations. Specifically, alignment of the striatum-DA weights to the DA-striatum weights was quantified, at every trial, by the correlation coefficient between the elements of **W**_SD_ and the elements of **C**_DS_. Alignment of the cortex(RNN)-striatum-DA connections to the DA-cortex(RNN) weights was quantified, at every trial, by the correlation coefficient between the elements of **W**_CS_**W**_SD_ and the elements of **C**_DC_. Correlation coefficient was calculated by corrcoef function of MATLAB.

Standard error of the mean (SEM) shown in the figures was approximated by SD (standard deviation)/√*N* (number of samples). Simulations were conducted by using MATLAB, and pseudo-random numbers were implemented by using rand, randn, and randperm functions. The codes for simulations and analyses will be made available at GitHub upon acceptance and publication of this manuscript at a journal.

## Author contributions

KM designed and performed the research in discussion with AK. KM wrote the first draft, and AK and KM edited the draft.

## Acknowledgements

KM was supported by Grants-in-Aid for Scientific Research 25H02594 and 23K27985 from Japan Society for the Promotion of Science (JSPS). AK acknowledges partial funding from the strategic research area StratNeuro.

## References

1. Montague, P.R., Dayan, P. & Sejnowski, T.J. A framework for mesencephalic dopamine systems based on predictive Hebbian learning. J Neurosci 16, 1936–1947 (1996).

2. Schultz, W., Dayan, P. & Montague, P.R. A neural substrate of prediction and reward. Science 275, 1593–1599 (1997).

3. Reynolds, J.N., Hyland, B.I. & Wickens, J.R. A cellular mechanism of reward-related learning. Nature 413, 67–70 (2001).

4. Hamid, A.A., et al. Mesolimbic dopamine signals the value of work. Nat Neurosci 19, 117–126 (2016).

5. de Jong, J.W., Liang, Y., Verharen, J.P.H., Fraser, K.M. & Lammel, S. State and rate-of-change encoding in parallel mesoaccumbal dopamine pathways. Nat Neurosci 27, 309–318 (2024).

6. Jeong, H., et al. Mesolimbic dopamine release conveys causal associations. Science 378, eabq6740 (2022).

7. Gershman, S.J., et al. Explaining dopamine through prediction errors and beyond. Nat Neurosci 27, 1645–1655 (2024).

8. Qian, L., et al. Prospective contingency explains behavior and dopamine signals during associative learning. Nat Neurosci 28, 1280–1292 (2025).

9. Kato, A. & Morita, K. Striatal Gradient in Value-Decay Explains Regional Differences in Dopamine Patterns and Reinforcement Learning Computations. J Neurosci e0170252025 (2025).

10. Takahashi, Y.K., et al. Dopamine Neurons Respond to Errors in the Prediction of Sensory Features of Expected Rewards. Neuron 95, 1395–1405.e1393 (2017).

11. Howard, J.D. & Kahnt, T. Identity prediction errors in the human midbrain update reward-identity expectations in the orbitofrontal cortex. Nat Commun 9, 1611 (2018).

12. Kahnt, T. & Schoenbaum, G. The curious case of dopaminergic prediction errors and learning associative information beyond value. Nat Rev Neurosci 26, 169–178 (2025).

13. Cools, R. & D’Esposito, M. Inverted-U-shaped dopamine actions on human working memory and cognitive control. Biol Psychiatry 69, e113–125 (2011).

14. Puig, M.V., Rose, J., Schmidt, R. & Freund, N. Dopamine modulation of learning and memory in the prefrontal cortex: insights from studies in primates, rodents, and birds. Front Neural Circuits 8, 93 (2014).

15. Ott, T. & Nieder, A. Dopamine and Cognitive Control in Prefrontal Cortex. Trends Cogn Sci 23, 213–234 (2019).

16. Puig, M.V. & Miller, E.K. The role of prefrontal dopamine D1 receptors in the neural mechanisms of associative learning. Neuron 74, 874–886 (2012).

17. D’Ardenne, K., et al. Role of prefrontal cortex and the midbrain dopamine system in working memory updating. Proc Natl Acad Sci U S A 109, 19900–19909 (2012).

18. Angrist, B.M. & Gershon, S. The phenomenology of experimentally induced amphetamine psychosis--preliminary observations. Biol Psychiatry 2, 95–107 (1970).

19. Kaar, S.J., Natesan, S., McCutcheon, R. & Howes, O.D. Antipsychotics: Mechanisms underlying clinical response and side-effects and novel treatment approaches based on pathophysiology. Neuropharmacology 172, 107704 (2020).

20. Nour, M.M., Liu, Y., El-Gaby, M., McCutcheon, R.A. & Dolan, R.J. Cognitive maps and schizophrenia. Trends Cogn Sci 29, 184–200 (2025).

21. Marder, S.R. & Galderisi, S. The current conceptualization of negative symptoms in schizophrenia. World Psychiatry 16, 14–24 (2017).

22. Marder, S.R. & Umbricht, D. Negative symptoms in schizophrenia: Newly emerging measurements, pathways, and treatments. Schizophr Res 258, 71–77 (2023).

23. Strauss, G.P., Waltz, J.A. & Gold, J.M. A review of reward processing and motivational impairment in schizophrenia. Schizophr Bull 40 Suppl 2, S107–116 (2014).

24. Gold, J.M., Waltz, J.A., Prentice, K.J., Morris, S.E. & Heerey, E.A. Reward processing in schizophrenia: a deficit in the representation of value. Schizophr Bull 34, 835–847 (2008).

25. Huys, Q.J.M., Browning, M., Paulus, M.P. & Frank, M.J. Advances in the computational understanding of mental illness. Neuropsychopharmacology 46, 3–19 (2021).

26. Millard, S.J., Bearden, C.E., Karlsgodt, K.H. & Sharpe, M.J. The prediction-error hypothesis of schizophrenia: new data point to circuit-specific changes in dopamine activity. Neuropsychopharmacology 47, 628–640 (2022).

27. Morris, R.W., Cyrzon, C., Green, M.J., Le Pelley, M.E. & Balleine, B.W. Impairments in action-outcome learning in schizophrenia. Transl Psychiatry 8, 54 (2018).

28. Heerey, E.A., Bell-Warren, K.R. & Gold, J.M. Decision-making impairments in the context of intact reward sensitivity in schizophrenia. Biol Psychiatry 64, 62–69 (2008).

29. Collins, A.G., Brown, J.K., Gold, J.M., Waltz, J.A. & Frank, M.J. Working memory contributions to reinforcement learning impairments in schizophrenia. J Neurosci 34, 13747–13756 (2014).

30. Collins, A.G.E., Albrecht, M.A., Waltz, J.A., Gold, J.M. & Frank, M.J. Interactions Among Working Memory, Reinforcement Learning, and Effort in Value-Based Choice: A New Paradigm and Selective Deficits in Schizophrenia. Biol Psychiatry 82, 431–439 (2017).

31. Robison, A.J., Thakkar, K.N. & Diwadkar, V.A. Cognition and Reward Circuits in Schizophrenia: Synergistic, Not Separate. Biol Psychiatry 87, 204–214 (2020).

32. Howes, O.D. & Shatalina, E. Integrating the Neurodevelopmental and Dopamine Hypotheses of Schizophrenia and the Role of Cortical Excitation-Inhibition Balance. Biol Psychiatry 92, 501–513 (2022).

33. Lewis, D.A., Hashimoto, T. & Volk, D.W. Cortical inhibitory neurons and schizophrenia. Nat Rev Neurosci 6, 312–324 (2005).

34. Insel, T.R. Rethinking schizophrenia. Nature 468, 187–193 (2010).

35. Grace, A.A. Dysregulation of the dopamine system in the pathophysiology of schizophrenia and depression. Nat Rev Neurosci 17, 524–532 (2016).

36. Millidge, B., Song, Y., Lak, A., Walton, M.E. & Bogacz, R. Reward Bases: A simple mechanism for adaptive acquisition of multiple reward types. PLoS Comput Biol 20, e1012580 (2024).

37. Collins, A.G. & Frank, M.J. Opponent actor learning (OpAL): modeling interactive effects of striatal dopamine on reinforcement learning and choice incentive. Psychol Rev 121, 337–366 (2014).

38. Keramati, M. & Gutkin, B. Homeostatic reinforcement learning for integrating reward collection and physiological stability. Elife 3, e04811 (2014).

39. Möller, M. & Bogacz, R. Learning the payoffs and costs of actions. PLoS Comput Biol 15, e1006285 (2019).

40. Wärnberg, E. & Kumar, A. Feasibility of dopamine as a vector-valued feedback signal in the basal ganglia. Proc Natl Acad Sci U S A 120, e2221994120 (2023).

41. Tsurumi, T., Kato, A., Kumar, A. & Morita, K. Online reinforcement learning of state representation in recurrent network: the power of random feedback and biological constraints. eLife https://elifesciences.org/reviewed-preprints/104101 (2025).

42. Hennig, J.A., et al. Emergence of belief-like representations through reinforcement learning. PLoS Comput Biol 19, e1011067 (2023).

43. Doya, K. Complementary roles of basal ganglia and cerebellum in learning and motor control. Curr Opin Neurobiol 10, 732–739 (2000).

44. Lillicrap, T.P., Cownden, D., Tweed, D.B. & Akerman, C.J. Random synaptic feedback weights support error backpropagation for deep learning. Nat Commun 7, 13276 (2016).

45. Murray, J.M. Local online learning in recurrent networks with random feedback. Elife 8, e43299 (2019).

46. Bellec, G., et al. A solution to the learning dilemma for recurrent networks of spiking neurons. Nat Commun 11, 3625 (2020).

47. Niv, Y., Daw, N.D., Joel, D. & Dayan, P. Tonic dopamine: opportunity costs and the control of response vigor. Psychopharmacology (Berl) 191, 507–520 (2007).

48. Rajan, K. & Abbott, L.F. Eigenvalue spectra of random matrices for neural networks. Phys Rev Lett 97, 188104 (2006).

49. Frank, M.J., Seeberger, L.C. & O’reilly, R.C. By carrot or by stick: cognitive reinforcement learning in parkinsonism. Science 306, 1940–1943 (2004).

50. Pinto, S.R. & Uchida, N. Tonic dopamine and biases in value learning linked through a 1 biologically inspired reinforcement learning model. bioRxiv 10.1101/2023.11.10.566580 (2023).

51. Elman, I., Borsook, D. & Lukas, S.E. Food intake and reward mechanisms in patients with schizophrenia: implications for metabolic disturbances and treatment with second-generation antipsychotic agents. Neuropsychopharmacology 31, 2091–2120 (2006).

52. Greenstreet, F., et al. Dopaminergic action prediction errors serve as a value-free teaching signal. Nature 643, 1333–1342 (2025).

53. Valjent, E. & Gangarossa, G. The Tail of the Striatum: From Anatomy to Connectivity and Function. Trends Neurosci 44, 203–214 (2021).

54. Jiang, H. & Kim, H.F. Anatomical Inputs From the Sensory and Value Structures to the Tail of the Rat Striatum. Front Neuroanat 12, 30 (2018).

55. Ogata, S., Miyamoto, Y., Shigematsu, N., Esumi, S. & Fukuda, T. The Tail of the Mouse Striatum Contains a Novel Large Type of GABAergic Neuron Incorporated in a Unique Disinhibitory Pathway That Relays Auditory Signals to Subcortical Nuclei. J Neurosci 42, 8078–8094 (2022).

56. Mueser, K.T., Bellack, A.S. & Brady, E.U. Hallucinations in schizophrenia. Acta Psychiatr Scand 82, 26–29 (1990).

57. Menegas, W., Babayan, B.M., Uchida, N. & Watabe-Uchida, M. Opposite initialization to novel cues in dopamine signaling in ventral and posterior striatum in mice. Elife 6 (2017).

58. Menegas, W., Akiti, K., Amo, R., Uchida, N. & Watabe-Uchida, M. Dopamine neurons projecting to the posterior striatum reinforce avoidance of threatening stimuli. Nat Neurosci 21, 1421–1430 (2018).

59. Akiti, K., et al. Striatal dopamine explains novelty-induced behavioral dynamics and individual variability in threat prediction. Neuron 110, 3789–3804.e3789 (2022).

60. Tsutsui-Kimura, I., et al. Dopamine in the tail of the striatum facilitates avoidance in threat-reward conflicts. Nat Neurosci 28, 795–810 (2025).

61. Krystal, J.H., et al. Subanesthetic effects of the noncompetitive NMDA antagonist, ketamine, in humans. Psychotomimetic, perceptual, cognitive, and neuroendocrine responses. Arch Gen Psychiatry 51, 199–214 (1994).

62. Jackson, M.E., Homayoun, H. & Moghaddam, B. NMDA receptor hypofunction produces concomitant firing rate potentiation and burst activity reduction in the prefrontal cortex. Proc Natl Acad Sci U S A 101, 8467–8472 (2004).

63. Murray, R.M. & Lewis, S.W. Is schizophrenia a neurodevelopmental disorder? Br Med J (Clin Res Ed) 295, 681–682 (1987).

64. Marenco, S. & Weinberger, D.R. The neurodevelopmental hypothesis of schizophrenia: following a trail of evidence from cradle to grave. Dev Psychopathol 12, 501–527 (2000).

65. Murray, R.M., Bhavsar, V., Tripoli, G. & Howes, O. 30 Years on: How the Neurodevelopmental Hypothesis of Schizophrenia Morphed Into the Developmental Risk Factor Model of Psychosis. Schizophr Bull 43, 1190–1196 (2017).

66. Jardri, R. & Denève, S. Circular inferences in schizophrenia. Brain 136, 3227–3241 (2013).

67. Murray, J.D., et al. Linking microcircuit dysfunction to cognitive impairment: effects of disinhibition associated with schizophrenia in a cortical working memory model. Cereb Cortex 24, 859–872 (2014).

68. Jardri, R., Duverne, S., Litvinova, A.S. & Denève, S. Experimental evidence for circular inference in schizophrenia. Nat Commun 8, 14218 (2017).

69. Lanillos, P., et al. A review on neural network models of schizophrenia and autism spectrum disorder. Neural Netw 122, 338–363 (2020).

70. Calvin, O.L. & Redish, A.D. Global disruption in excitation-inhibition balance can cause localized network dysfunction and Schizophrenia-like context-integration deficits. PLoS Comput Biol 17, e1008985 (2021).

71. Lam, N.H., et al. Effects of Altered Excitation-Inhibition Balance on Decision Making in a Cortical Circuit Model. J Neurosci 42, 1035–1053 (2022).

72. Langdon, C., Genkin, M. & Engel, T.A. A unifying perspective on neural manifolds and circuits for cognition. Nat Rev Neurosci 24, 363–377 (2023).

73. Hennequin, G., Agnes, E.J. & Vogels, T.P. Inhibitory Plasticity: Balance, Control, and Codependence. Annu Rev Neurosci 40, 557–579 (2017).

74. Kato, A., Kunisato, Y., Katahira, K., Okimura, T. & Yamashita, Y. Computational Psychiatry Research Map (CPSYMAP): A New Database for Visualizing Research Papers. Front Psychiatry 11, 578706 (2020).

75. Radulescu, A. & Niv, Y. State representation in mental illness. Curr Opin Neurobiol 55, 160–166 (2019).

76. Dalgleish, T., Black, M., Johnston, D. & Bevan, A. Transdiagnostic approaches to mental health problems: Current status and future directions. J Consult Clin Psychol 88, 179–195 (2020).

77. Redish, A.D. Addiction as a computational process gone awry. Science 306, 1944–1947 (2004).

78. Redish, A.D., Jensen, S. & Johnson, A. A unified framework for addiction: vulnerabilities in the decision process. Behav Brain Sci 31, 415–437; discussion 437-487 (2008).

79. Keiflin, R. & Janak, P.H. Dopamine Prediction Errors in Reward Learning and Addiction: From Theory to Neural Circuitry. Neuron 88, 247–263 (2015).

80. Speers, L.J., Schmidt, R. & Bilkey, D.K. Aberrant Phase Precession of Lateral Septal Cells in a Maternal Immune Activation Model of Schizophrenia Risk May Disrupt the Integration of Location with Reward. J Neurosci 42, 4187–4201 (2022).

81. Keidel, K., Murawski, C., Pantelis, C. & Ettinger, U. The Relationship Between Schizotypal Personality Traits and Temporal Discounting: The Role of the Date/Delay Effect. Schizophr Bull 51, S64–S73 (2025).

82. Heerey, E.A., Robinson, B.M., McMahon, R.P. & Gold, J.M. Delay discounting in schizophrenia. Cogn Neuropsychiatry 12, 213–221 (2007).

83. Wang, L., et al. Increased delayed reward during intertemporal decision-making in schizophrenic patients and their unaffected siblings. Psychiatry Res 262, 246–253 (2018).

84. Wang, Y., et al. Heterogeneity in the pyramidal network of the medial prefrontal cortex. Nat Neurosci 9, 534–542 (2006).

85. Mongillo, G., Barak, O. & Tsodyks, M. Synaptic theory of working memory. Science 319, 1543–1546 (2008).

86. Morishima, M., Morita, K., Kubota, Y. & Kawaguchi, Y. Highly differentiated projection-specific cortical subnetworks. J Neurosci 31, 10380–10391 (2011).

87. Bittner, K.C., Milstein, A.D., Grienberger, C., Romani, S. & Magee, J.C. Behavioral time scale synaptic plasticity underlies CA1 place fields. Science 357, 1033–1036 (2017).

88. Caya-Bissonnette, L., Naud, R. & Béïque, J.-C. Cellular Substrate of Eligibility Traces. bioRxiv 10.1101/2023.06.29.547097 (2023).

